# Scalable proxiloids enable human-relevant assessment of kidney proximal tubule toxicity

**DOI:** 10.64898/2026.08.31.748260

**Authors:** Sam Biermans, Dara N. van Heugten, M. Cristina Avramut, Roos-Sanne F. Verkerk, Rianne Y. van Nieuwland, Roan H. van Scheppingen, Linda van den Berk, Cathelijne W. van den Berg, Bob van de Water, Ton J. Rabelink, Mazène Hochane, H. Siebe Spijker

## Abstract

Drug-induced injury to the human proximal tubule (PT) is a leading cause of acute kidney injury and drug attrition, yet remains difficult to predict preclinically. PT toxicity arises from the coupling of transporter-mediated xenobiotic accumulation and high oxidative metabolic demand. Current models lack key aspects of PT physiology or are difficult to scale for toxicity testing. New Approach Methodologies (NAMs) address this challenge through human-relevant *in vitro* systems. Here we introduce proxiloids, a scalable suspension-based human induced pluripotent stem cell differentiation strategy. Within 14 days, proxiloids form lumenized, polarized tubular organoids enriched for PT identity, with functional transport and oxidative metabolic competence. Proxiloids are compatible with genetically encoded reporters and standard multiwell assays, enabling detection of defined stress responses. They recapitulate aminoglycoside nephrotoxicity with greater sensitivity than matched two-dimensional cultures and detect adefovir-induced mitochondrial toxicity not predicted in rodents. Together, proxiloids provide a scalable, human-relevant NAM for PT nephrotoxicity assessment.

**Teaser:** Preserved proximal tubule architecture enables human-relevant modeling of drug-induced kidney injury.

## INTRODUCTION

The development of new therapeutics requires careful assessment of potential toxicities throughout preclinical and clinical testing. Drug-induced nephrotoxicity remains a major safety concern, contributing to acute kidney injury, dose limitation, treatment discontinuation, and post-marketing drug withdrawal (*1, 2*). Within the kidney, injury predominantly involves the proximal tubule (PT), where toxicity arises from a combination of transporter-mediated xenobiotic uptake and high oxidative metabolic demand (*3*). PT cells express high levels of xenobiotic transporters that concentrate compounds intracellularly, while relying almost exclusively on oxidative phosphorylation to sustain function (*3, 4*). This coupling of active accumulation and high metabolic demand renders the PT particularly susceptible to compounds that exploit transporter pathways or impair mitochondrial function, enabling toxicity at concentrations well-tolerated by other cell types (*5*).

These same biological characteristics also make PT toxicity particularly difficult to model. Transporter activity, mitochondrial sensitivity, and epithelial polarity are highly species- and context-dependent, and together define the physiological state through which PT injury occurs (*6–8*). Consequently, animal models often fail to accurately recapitulate human PT injury, while *in vitro* systems frequently lack the physiological features required to elicit clinically relevant toxic responses. This limitation is exemplified by adefovir dipivoxil, which passed standard rodent toxicology testing yet caused PT nephrotoxicity in patients through Organic Anion Transporter (OAT)-mediated intracellular accumulation and mitochondrial dysfunction (*9–11*). Such cases have contributed to growing regulatory and scientific recognition that human-relevant models are needed to bridge the gap between preclinical testing and clinical outcomes.

These efforts have accelerated the development of New Approach Methodologies (NAMs). Importantly, the FDA Modernization Act 2.0 removed the statutory requirement for animal studies prior to Investigational New Drug applications and explicitly encouraged the use of NAMs as scientifically rigorous complements to animal-based testing (*12, 13*). For a PT toxicity model, biological relevance depends on 1) reproducing key human PT features, including transporter-mediated uptake, epithelial polarization, and oxidative metabolism; 2) maintaining these features to enable mechanistically interpretable injury assessment that reflects clinically observed toxicity; and 3) remaining compatible with scalable, routine experimental workflows (*13*).

Existing human *in vitro* PT models meet these requirements only in part. Two-dimensional (2D) cultures, including HK-2, RPTEC/TERT1, conditionally immortalized PT cells, and human induced pluripotent stem cell (hiPSC)-derived monolayers, offer scalability but incompletely preserve tubular architecture and lack fully developed apical-basolateral polarity (*14–17*). The absence of tubular architecture disrupts the spatial organization of transporters and the metabolic specialization that underlie physiological injury responses (*18*). Microphysiological systems (MPS) improve structural fidelity and enable controlled transport measurements (*19–21*), but are constrained by low throughput, manufacturing complexity, and high costs. hiPSC-derived kidney organoids provide greater three-dimensional (3D) nephron-like structures and represent the most developmentally complete *in vitro* kidney model (*22–25*), yet have limited PT representation and exhibit variability, extended culture times, and poor scalability (*26, 27*). Thus, existing models each capture different aspects of PT physiology, but few combine physiological relevance, PT specificity, scalability, and experimental accessibility in a single platform.

Here, we introduce proxiloids, a suspension-based hiPSC differentiation strategy designed to combine PT-specific biology with scalable experimental workflows. By adapting a validated 2D PT differentiation protocol (*17*) to scalable suspension culture, proxiloids self-organize into lumenized, polarized tubular organoids enriched for PT identity and exhibit functional transporter activity and oxidative metabolic competence. The 14-day workflow generates hundreds of organoids with low within-batch morphological variability and reproducible transcriptional identity across hiPSC lines. Proxiloids are compatible with genetically encoded reporters and standard multiwell assays, enabling detection of defined cellular stress responses at bulk and cellular resolution. We further show that proxiloids recapitulate aminoglycoside nephrotoxicity with greater sensitivity than matched 2D cultures and detect adefovir-induced mitochondrial toxicity not predicted in rodents. Together, these findings establish proxiloids as a scalable, human-relevant system for mechanism-resolved PT nephrotoxicity assessment.

## RESULTS

### Proxiloids combine tubular architecture, transporter function, and mitochondrial metabolism

To establish a scalable PT organoid model that incorporates key architectural features of the native tubule, we developed a 3D implementation of a previously validated protocol for 2D PT-like cells (*17*). This strategy combines directed PT specification with 3D tissue organization in suspension culture. hiPSCs were seeded in a microwell culture system to control homogenous aggregation, generating 1,500 embryoid bodies (EBs) per well (Fig. 1A). Upon transfer to suspension, EBs underwent a uniform morphological progression, exhibiting early epithelial organization by day 3 and forming structured tubular organoids, hereafter termed proxiloids, by day 14 (Fig. 1B).

**Fig. 1.**
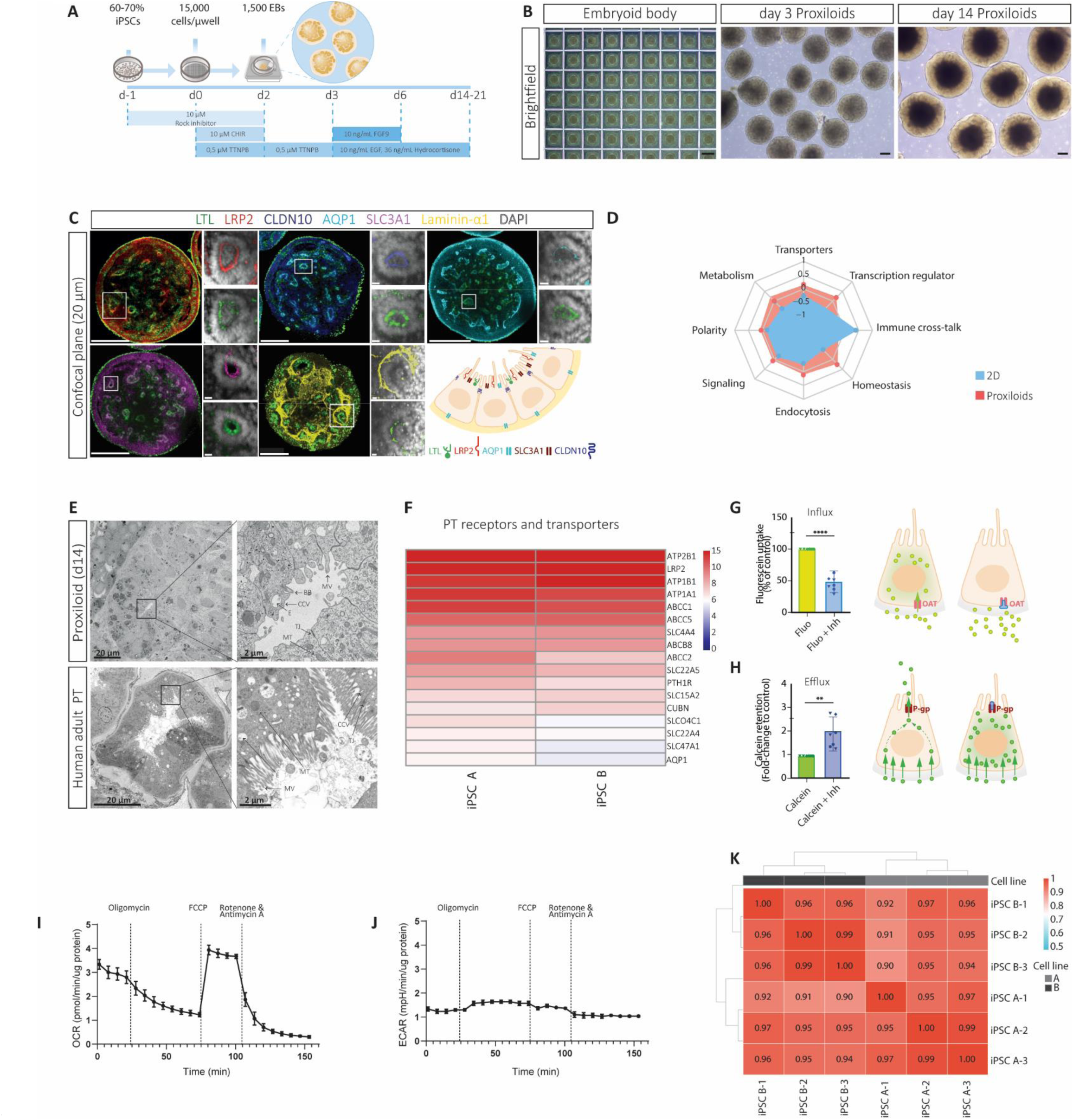
Proxiloids integrate tubular architecture, transporter function, and mitochondrial metabolism. **(A)** Schematic of microwell-based suspension differentiation strategy for proxiloid generation. **(B)** Brightfield images at day −1 (scale bar, 500 µm), and days 3 and 14 (scale bars, 200 µm). **(C)** Confocal immunofluorescence showing apical markers (LRP2, AQP1, SLC3A1, LTL), basolateral laminin-α1, and tight junction protein claudin-10 (CLDN10); DAPI labels nuclei. Scale bars, 100 µm; zoom, 10 µm. **D)** PT signature category scores, comparing proxiloids and 2D monolayers; values are shown as z-scored mean expression. **(E)** TEM of proxiloids (top) and human proximal tubule (bottom). MV, microvilli; TJ, tight junction; M, mitochondria; E, endosome; CCV, clathrin-coated vesicle; BB, basal body. **(F)** Heatmap of variance-stabilized transformed (VST) expression levels of xenobiotic transporters from bulk RNA-sequencing averaged across two hiPSC lines (n=3 per line). **(G)** Fluorescein (1 µM) uptake ± probenecid (500 µM); Inh, inhibitor; mean ± SD, n=3 lines; ****P < 0.0001, paired t-test. **(H)** Calcein retention (0.25 µM) ± PSC-833 (5 µM); Inh, inhibitor; mean ± SD, n=3 lines; **P < 0.01, paired t-test. **(I-J)** OCR traces following mitochondrial stress test **(I)** and corresponding ECAR measurements **(J)**; data are shown as mean ± SEM. **(K)** Genome-wide Spearman correlation between two hiPSC lines (n=3 biological replicates per line).

Immunofluorescence imaging demonstrated that proxiloids contain polarized epithelial tubules with clear apical-basolateral organization (Fig. 1C). PT markers megalin (*LRP2*), aquaporin-1 (*AQP1*), rBAT (*SLC3A1*), and Lotus tetragonolobus lectin (LTL) localized to the apical luminal surface, while laminin-α1 (*LAMA1*) and claudin-10 (*CLDN10*) defined the basolateral compartment and tight junctions, respectively. 3D reconstruction revealed numerous extended, lumenized tubular networks throughout the organoid (Fig. S1A, Movie S1).

To assess PT identity at the transcriptional level, we performed bulk RNA sequencing on day 14 proxiloids from two hiPSC lines. Comparison with a previously published dataset of 2D PT-like cells (*17*) showed higher expression of a curated PT signature (*28*) in proxiloids across both lines (Fig. S1B), with enrichment across key functional categories central to PT physiology, including transport, polarity, and metabolism (Fig. 1D, Fig. S1C, Data S1). Consistent with enhanced PT enrichment, markers of non-PT nephron segments had lower expression in proxiloids than in the published 2D cultures by bulk RNA sequencing (Fig. S1D) and were absent by immunostaining (Fig. S1E). Stromal-associated genes, including *COL1A2*, *COL3A1*, and *COL6A3*, as well as lateral plate mesoderm markers, were also reduced relative to 2D cultures. Muscle markers were absent or minimally expressed, while a neural population remained detectable in both 2D cultures and proxiloids, reflected by expression of *STMN2*, *ELAVL3*, and *MAP2* (Fig S1D), consistent with off-target populations previously reported in kidney organoids (26).

Ultrastructural analysis by transmission electron microscopy confirmed epithelial maturation, including luminal organization, microvilli forming a brush border, tight junctions, and elongated mitochondria (Fig. 1E). Additional features such as endosomal structures, clathrin-coated vesicles, and primary cilia were observed, consistent with active epithelial and endocytic function. Proxiloids reproduced the principal ultrastructural features of native human PT, although luminal diameter, mitochondrial density, and brush border length remained reduced, consistent with a maturing *in vitro* state. By day 21, proxiloids exhibited expanded lumens while maintaining polarized epithelial organization (Fig. S1F).

Because selective transporter-mediated uptake drives intracellular accumulation of many nephrotoxic compounds in PT cells, we assessed transporter expression and function. Bulk RNA sequencing demonstrated expression of a broad repertoire of PT transport systems involved in xenobiotic handling (Fig. 1F). To confirm that transporter expression translated into active, inhibitor-sensitive transport, we performed functional uptake and efflux assays. Organic anion uptake measured by fluorescein accumulation was reduced by 51% (SEM ± 5.6%) upon OAT inhibition with probenecid, while intracellular calcein retention increased by 97% (SEM ± 24.4%) upon P-glycoprotein inhibition with PSC-833 (Fig. 1G–H), demonstrating functional basolateral uptake and apical efflux.

Given the role of oxidative metabolism in PT physiology and vulnerability to injury, we assessed bioenergetic function. Proxiloids exhibited high basal oxygen consumption (OCR) and low extracellular acidification (ECAR), yielding an OCR to ECAR ratio of ∼3:1 and indicating predominant reliance on oxidative phosphorylation (Fig. 1I-J). Consistent with this, mitochondrial stress testing revealed a marked increase in OCR following FCCP addition, indicating substantial spare respiratory capacity. ECAR remained comparatively low, suggesting limited glycolytic compensation and resembling the metabolic behavior of native human PT cells.

Lastly, we investigated the reproducibility and scalability of the platform. Proxiloids formed with consistent morphology across differentiations in three independent hiPSC lines, with minimal size variability (coefficient of variation (CV) <6% within batches, <16% between batches; Fig. S1G-H). This morphological consistency extended to transcriptional identity, where genome-wide Spearman correlations exceeded 0.90 both within and across cell lines (Fig. 1K), demonstrating stable and cell line-independent transcriptional identity. Together with the generation of hundreds of organoids per differentiation, these findings establish a 14-day platform that combines PT-enriched tubular architecture, functional transport and oxidative metabolism with reproducibility and scale suitable for routine multiwell experiments.

### Proxiloids enable detection of a canonical DNA-damage response

Next, we asked whether proxiloids can detect a well-defined cellular stress response. Proxiloids were exposed to etoposide (30 µM, 24 h), a topoisomerase II inhibitor that leads to DNA double-strand breaks and activates canonical p53-dependent signaling, providing a mechanistically defined, non–PT-specific stress stimulus (*29, 30*).

Bulk RNA sequencing revealed extensive transcriptional remodeling, with 2,386 genes significantly altered relative to controls (adjusted p < 0.01, |log2FC| > 1.5; Fig. 2A). Among the most strongly upregulated transcripts were core p53 targets, including *CDKN1A* (p21), *MDM2*, and *GADD45A*, as well as pro-apoptotic mediators *BAX*, *BBC3* (PUMA), and *PMAIP1* (NOXA), consistent with activation of a canonical DNA-damage response.

**Fig. 2.**
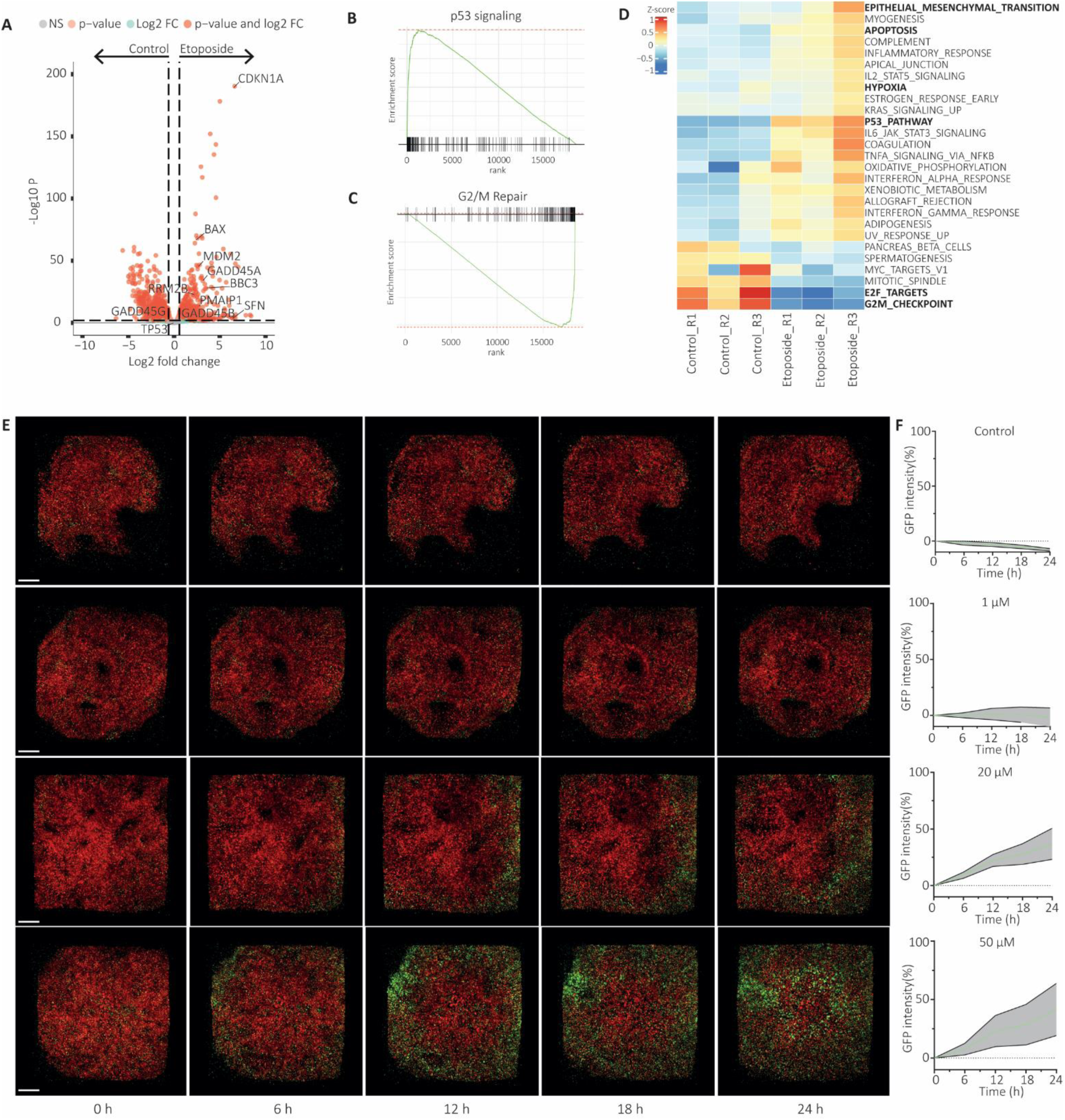
Proxiloids enable detection of a canonical DNA-damage response. **(A)** Volcano plot of etoposide versus control (30 µM, 24 h; n = 3). Thresholds: adjusted P value (Padj) = 0.01 and |log2FC| = 1.5 (dashed lines). Selected p53 and apoptosis-related genes are labeled; 2,386 differentially expressed genes (DEGs) identified. **(B–C)** Gene set enrichment analysis (GSEA) of Hallmark pathways: **(B)** p53 signaling (NES = 3.03, Padj < 0.001) and **(C)** G2/M checkpoint (NES = −2.96, Padj < 0.001). **(D)** Hallmark pathway activity (mean z-score across pathway genes) in control and etoposide samples; significant pathways (Padj < 0.05) shown. **(E)** Time-lapse confocal imaging of p21-GFP reporter proxiloids from 0–24 h following vehicle or etoposide treatment; scale bar, 100 µm. **(F)** Quantification of nuclear GFP fluorescence expressed as percentage of baseline over 24 h across concentrations; mean ± SEM, n = 3.

Gene set enrichment analysis confirmed strong induction of p53 signaling (Normalized Enrichment Score (NES) = 3.03) and apoptosis pathways (NES = 2.12), coupled to suppression of proliferative programs (Fig. 2B–C). Etoposide markedly downregulated E2F targets (NES = −2.82), G2/M checkpoint genes (NES = −2.96), and MYC-associated transcriptional signatures (NES = −1.77), consistent with cell cycle arrest and a shift toward stress and repair states. Pathway-level scoring further demonstrated activation of apoptosis, hypoxia, and epithelial remodeling pathways (Fig. 2D). Together, these data demonstrate that proxiloids can detect a robust, canonical transcriptional response to a mechanistically defined DNA-damaging stimulus.

To assess whether this response could be monitored dynamically, we generated proxiloids from a p21-GFP reporter hiPSC line. In this line, GFP expression reports induction of *CDKN1A* (p21), a canonical downstream target of p53 and marker of cellular stress and DNA damage responses. Time-lapse imaging revealed a progressive, dose-dependent increase in nuclear GFP signal following etoposide exposure, detectable within hours and increasing over 24 hours (Fig. 2E). Higher concentrations accelerated response kinetics, while untreated controls remained stable (Fig. 2F). These data demonstrate that proxiloids are compatible with dynamic, reporter-based detection of cellular stress responses at cellular resolution.

### Gentamicin-induced proximal tubule injury is recapitulated in proxiloids

To assess PT-specific injury responses, proxiloids were exposed to the aminoglycoside antibiotic gentamicin, a nephrotoxicant that accumulates in PT cells via megalin-mediated endocytosis and induces mitochondrial dysfunction (*31*).

Exposure to gentamicin (4 mg/mL) impaired mitochondrial function after 24 and 48 hours of treatment. At 24 hours, modest reductions in basal and ATP-linked respiration were accompanied by significantly increased spare respiratory capacity, consistent with an early compensatory response (Fig. 3A, C-E). By 48 hours, gentamicin suppressed oxidative phosphorylation, with significant reductions in basal respiration, ATP production, and non-mitochondrial respiration (Fig. 3A, C-F). Extracellular acidification remained low across conditions, indicating limited glycolytic compensation and consistent with the predominantly oxidative metabolic phenotype of PT epithelium (Fig. 3B).

**Fig. 3.**
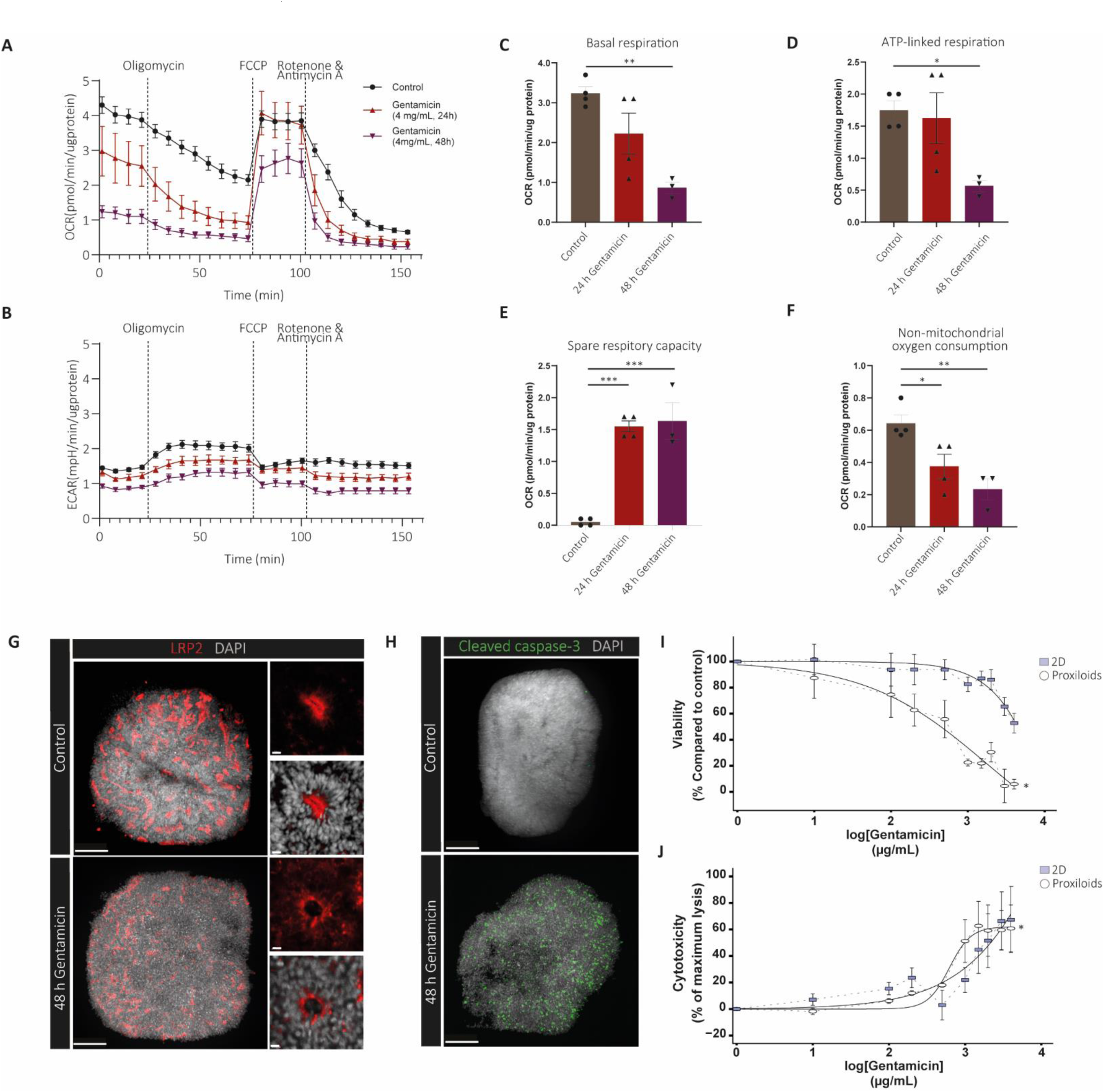
Proxiloids recapitulate gentamicin-induced injury and exhibit increased sensitivity. **(A–B)** Seahorse oxygen consumption rate (OCR) traces **(A)** and extracellular acidification rate (ECAR) **(B)** for control and gentamicin-treated proxiloids (4 mg/mL, 24 h and 48 h); injection points indicated. Data shown as mean ± SEM. **(C–F)** Quantification of mitochondrial respiration parameters normalized to control: **(C)** basal respiration, **(D)** ATP-linked respiration, **(E)** spare respiratory capacity, and **(F)** non-mitochondrial respiration; mean ± SEM, n = 3; *Padj < 0.05, **Padj < 0.01, ***Padj < 0.001, ordinary one-way ANOVA with Tukey’s multiple-comparison test. **(G)** LRP2 immunofluorescence, control versus 48 h gentamicin; scale bar, 100 µm; zoom, 10 µm. **(H)** Cleaved caspase-3 immunofluorescence, control versus 48 h gentamicin; DAPI labels nuclei; scale bar, 100 µm. **(I–J)** Dose-response comparison between proxiloids and matched 2D cultures. **(I)** WST-8 viability (EC50 923 µg/mL in proxiloids; no reliable EC50 in 2D, viability >50% across the tested range) and **(J)** LDH release (IC50 668 µg/mL in proxiloids vs 1,460 µg/mL in 2D). Data are shown as mean ± SEM, n = 3, *P < 0.05. Fitted with a four-parameter log-logistic model.

Consistent with PT injury, gentamicin exposure disrupted epithelial organization and activated cell death pathways. Immunostaining revealed loss of sharp apical LRP2 localization and fragmentation of tubular structures, accompanied by induction of cleaved caspase-3, consistent with activation of apoptotic signaling (Fig. 3G-H).

To benchmark gentamicin sensitivity against a conventional culture format, we generated matched 2D hiPSC-derived PT monolayers from the same parental hiPSC line using the previously established PT differentiation protocol (*17*). Dose-response analysis demonstrated concentration-dependent cytotoxicity in proxiloids, with an EC50 of 923 µg/mL for metabolic viability and an IC50 of 668 µg/mL for LDH release (Fig. 3I-J). In matched 2D hiPSC-derived PT monolayers, viability remained above 50% across the tested concentration range, precluding reliable estimation of an EC50, whereas the LDH IC50 was 1,460 µg/mL. Proxiloids therefore exhibited a 2.2-fold lower LDH IC50 than matched 2D cultures, indicating greater sensitivity to gentamicin-induced injury in the 3D culture format.

Together, these findings demonstrate that proxiloids reproduce key features of gentamicin-induced PT injury, including disruption of PT epithelial organization, mitochondrial dysfunction, and apoptotic signaling, and exhibit greater sensitivity to gentamicin-induced cytotoxicity than matched 2D cultures.

### Proxiloids model adefovir-induced mitochondrial nephrotoxicity

To assess whether proxiloids capture PT toxicity not detected in conventional preclinical animal models, we exposed proxiloids to adefovir, the active moiety of the prodrug adefovir dipivoxil that is associated with mitochondrial PT injury in patients (*9*). Adefovir was used directly to model renal epithelial exposure to the active drug. Proxiloids were exposed to 0.01-5 µM adefovir, a range spanning the reported patient Cmax (18 ng/mL, ∼0.07 µM)(*32*) and extending to concentrations potentially reached in the PT through OAT1-mediated tubular accumulation (*33*).

Exposure of proxiloids to adefovir disrupted mitochondrial integrity in a dose-dependent manner. Quantification of mitochondrial DNA (mtDNA) revealed a significant reduction in mtDNA copy number relative to nuclear DNA after seven days of repeated exposure (Fig. 4A).

**Fig.4.**
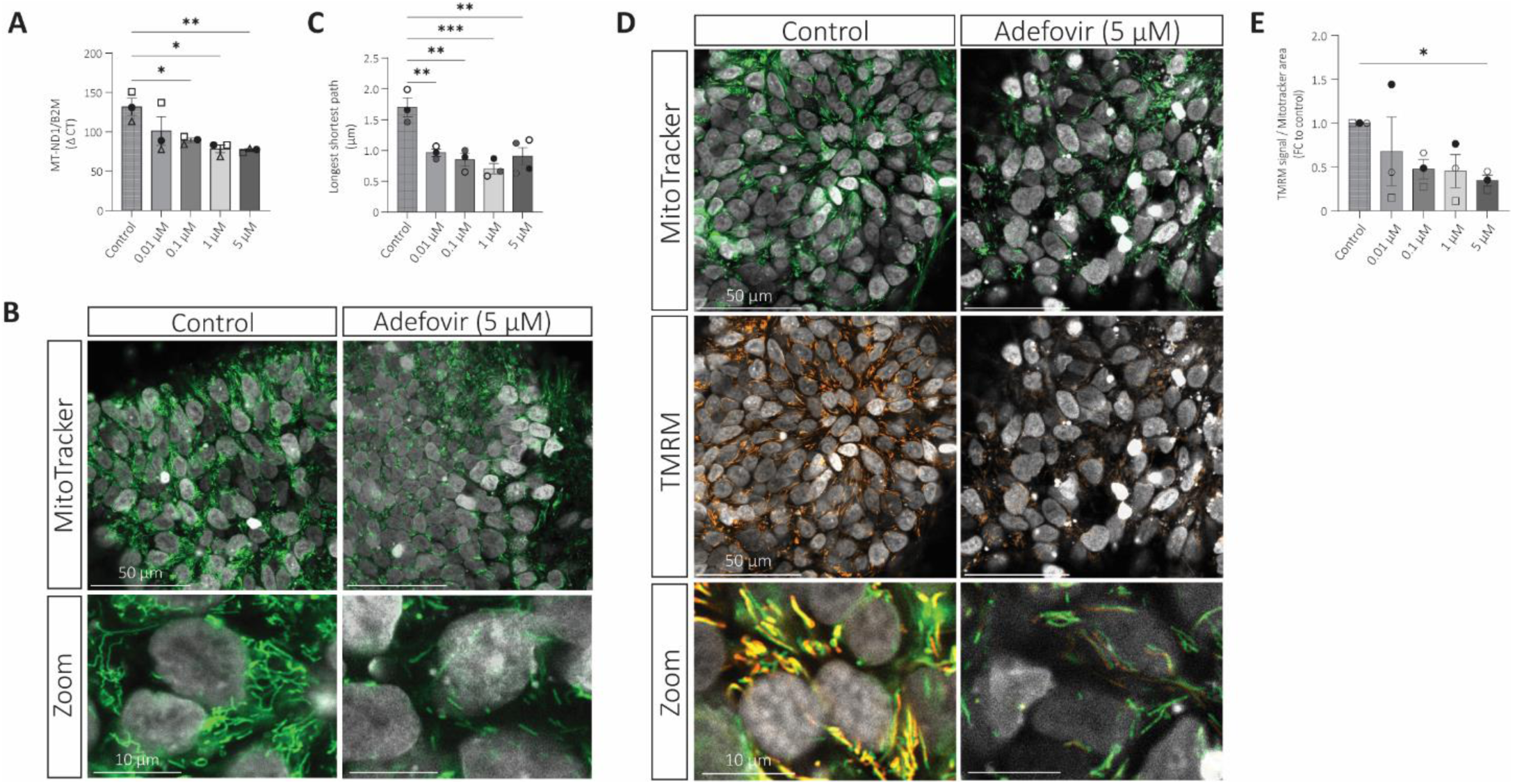
Proxiloids detect adefovir-induced mitochondrial nephrotoxicity at concentrations relevant to renal tubular exposure. **(A)** Mitochondrial DNA copy number measured at the MT-ND1/B2M ratio after 7 days of adefovir exposure across concentrations; mean ± SEM, n = 3; one-way ANOVA with Dunnett’s multiple-comparison test; *Padj < 0.05, **Padj < 0.01. **(B)** Mitotracker Green staining showing mitochondrial morphology in control- and adefovir-treated proxiloids following 3 days of exposure. **(C)** Quantification of mitochondrial network connectivity (longest shortest path) across adefovir concentrations; mean ± SEM, n = 3; one-way ANOVA with Dunnett’s multiple-comparison test; **Padj < 0.01, ***Padj < 0.001. **(D)** TMRM staining showing mitochondrial membrane potential in control- and adefovir-treated proxiloids following 3 days of exposure. **(E)** Quantification of TMRM fluorescence expressed as percentage of control across concentrations; mean ± SEM, n = 3; one-way ANOVA with Dunnett’s multiple-comparison test; *Padj < 0.05.

At an earlier timepoint of three days, high-resolution imaging revealed a shift from elongated, interconnected mitochondrial networks in controls to fragmented and punctate structures following adefovir exposure (Fig. 4B). Quantitative analysis confirmed a reduction in mitochondrial network connectivity, reflected by decreased longest shortest path measurements across conditions (Fig. 4C).

Loss of mitochondrial function was further reflected by reduced membrane polarization after three days. Combined Mitotracker and TMRM imaging, with TMRM reporting mitochondrial membrane potential, showed a clear reduction in TMRM signal intensity following adefovir exposure (Fig. 4D-E).

Together, these findings show that proxiloids reproduce key cellular features of adefovir-induced PT mitochondrial injury, including mtDNA depletion, mitochondrial network fragmentation, and loss of membrane potential.

## DISCUSSION

In the present study, we set out to address a practical limitation in modeling PT toxicity. Existing systems often require a trade-off between physiological relevance and experimental scalability. We therefore developed proxiloids, a PT-enriched hiPSC-derived organoid system that combines lumenized tubular architecture and epithelial polarity with functional transport, oxidative metabolism, and scalable suspension culture. Proxiloids were reproducibly generated across hiPSC lines, with low within-batch morphological variability and stable transcriptional identity, and were compatible with genetically encoded reporters and standard assay platforms. Functionally, they enabled detection of a canonical cellular stress response and recapitulated PT-specific injury, including increased sensitivity to gentamicin relative to matched 2D cultures and adefovir-induced mitochondrial toxicity not predicted in rodent models. These features position proxiloids as a complementary model between conventional PT monolayers, complex kidney organoids, and technically demanding MPS systems.

PT cells are known to progressively lose epithelial polarity, transporter organization, and metabolic specialization in conventional 2D culture, leading to reduced uptake-dependent injury responses (*34, 35*). Restoring a 3D environment has been shown to recover key aspects of PT physiology and enhance sensitivity to nephrotoxins (*18, 36*). Our data provide evidence that this structural context contributes to the functional response to gentamicin. Although the transcriptomic comparison between proxiloids and 2D PT-like cells was based on independently generated datasets, the gentamicin experiments directly compared proxiloids and 2D monolayers generated from the same differentiation protocol and cell source. Proxiloids exhibited an LDH IC50 of 668 µg/mL compared with 1,460 µg/mL in matched 2D cultures, while an EC50 for metabolic viability could not be reliably estimated in 2D because viability remained above 50% across the tested concentration range. Because these cultures shared a common cellular origin, the increased sensitivity in proxiloids is consistent with contributions from 3D epithelial organization, transporter activity, and metabolic state rather than lineage specification alone.

Proxiloids responded to mechanistically defined cellular insults. Gentamicin induced mitochondrial dysfunction and disruption of epithelial organization, consistent with megalin-mediated aminoglycoside accumulation and its role in oxidative stress and mitochondrial injury (*36*). In parallel, etoposide induced a transcriptional response consistent with its established mechanism, in which topoisomerase II inhibition and DNA double-strand breaks activate p53 signaling and cell-cycle arrest (*29, 37*). This response was independently monitored using a p21-GFP reporter line, which showed dynamic upregulation of p21 following etoposide exposure at cellular resolution. This demonstrates that proxiloids are suitable for longitudinal imaging and quantitative analysis of genetically encoded stress-response reporters. Combined with their scalability, these features provide a foundation for high-content drug screening and mechanistic studies that are difficult to perform in primary systems and challenging to scale in MPS. Since proxiloids are hiPSC-derived, this reporter-based approach can readily be extended to targeted perturbations and defined genetic backgrounds, further supporting applications in disease modeling.

The response to adefovir provides a particularly important example of the added value of a human PT model. Adefovir nephrotoxicity is linked to OAT-mediated uptake and intracellular accumulation in PT cells, followed by mitochondrial injury (*9, 11*). This may contribute to the limited predictive value of conventional rodent models, as human OAT1 has substantially greater affinity for adefovir than the rat ortholog (*6, 8*). Clinically, adefovir exposure following therapeutic dosing is in the submicromolar range, with reported plasma Cmax value of approximately 0.07 µM (*32*). Our lowest exposure, 0.01 µM, was below this range, while 0.1 µM approximated reported systemic exposure. Higher concentrations up to 5 µM were used to assess the consequences of increased extracellular exposure and potential transporter-mediated intracellular accumulation. Proxiloids reproduced adefovir-induced mtDNA depletion, mitochondrial fragmentation, and loss of membrane potential across this exposure range, demonstrating their ability to capture a human PT-specific mitochondrial toxicity phenotype not predicted in conventional preclinical models. This alignment between mechanism, measurable endpoints, and clinically observed toxicity supports the core biological-relevance criteria for human-relevant NAMs. Systematic benchmarking across compound classes will be needed to further define its predictive scope.

These properties position proxiloids as a complementary PT-focused model within the existing landscape of kidney *in vitro* systems. Conventional monolayer cultures provide accessible and scalable platforms but do not fully preserve the structural and metabolic features that contribute to PT-specific injury (*34*). Kidney organoids reproduce nephron-level complexity but contain heterogenous cell populations that complicate the assessment of PT-specific responses (*26, 27*). Primary tubuloids recover functional epithelium but are donor-dependent and less susceptible to genetic manipulation (*38*). MPS provide polarized transport and fluid flow but are technically demanding and difficult to scale (*19–21, 39*). Proxiloids instead combine PT enrichment and tubular organization with a 14-day differentiation workflow, reproducible production across hiPSC lines, genetic accessibility, and compatibility with standard multiwell assays. Their value therefore lies not in replacing existing models, but in providing a scalable PT-focused system that integrates structural and functional features relevant to nephrotoxicity.

Several aspects define the current scope of the platform. PT enrichment was supported by bulk transcriptomic, structural, and functional analyses. However, the degree to which individual PT subsegments are represented remains unknown and may influence applications focused on segment-specific toxicities (*40, 41*). Alongside the PT epithelium, proxiloids also contained off-target populations. A stromal population was present, indicated by expression of collagen genes *COL1A2*, *COL3A1*, and *COL6A3* (*42*). Stromal and mesenchymal cells have previously been shown to support epithelial polarization and tubular organization in kidney and other organoid systems (*43*). These populations may additionally influence toxicological responses through paracrine interactions, although their contribution to the injury phenotypes observed in proxiloids remains unclear. A neural population was also detected, indicated by expression of *STMN2*, *ELAVL3*, and *MAP2*. Neural differentiation is a recognized off-target outcome of hiPSC protocols, including kidney organoid differentiation, arguably reflecting the tendency of pluripotent cells toward neuroectoderm when lineage restriction is incomplete (*26, 44*). Both populations may arise from incompletely restricted progenitors during the early phase of differentiation, and reducing these off-target populations through more stringent intermediate mesoderm specification may be an important direction for further studies. In the present study, we focused on PT-intrinsic responses and incorporation of endothelial or immune compartments was not assessed. Their integration, for example through assembloid approaches, is a promising direction for modeling the multicellular contributions to nephrotoxicity *in vivo*.

Together, these findings show that preserving PT physiological function is an important determinant of toxic injury responses *in vitro* and establish proxiloids as a scalable, human-relevant platform for mechanistic study of PT injury. By combining PT-enriched tubular architecture, functional transport and oxidative metabolism with reproducible production and routine experimental accessibility, proxiloids provide a complementary approach for mechanism-resolved nephrotoxicity studies and the development of NAM-based strategies for renal safety assessment.

## MATERIALS AND METHODS

### hiPSC maintenance

hiPSC cultures were maintained in an undifferentiated state in mTeSR^TM^ Plus medium (StemCell Technologies, cat.no. 100-0276) on Growth Factor Reduced Matrigel-coated 6-well plates or T75 flasks (Corning, cat. no. 354230; Greiner, cat. no. 628161). We primarily used the commercially available iPSC0028 line (Sigma-Aldrich; iPSC A). An iPSC0028-HC3X-derived CDKN1A^GFP^-H2B^Cy5^ reporter line was generated by CRISPR/Cas9-mediated genome editing for toxicity reporter studies. The HC3X background contains doxycycline-inducible *HNF1A*, *FOXA3*, and *PROX1* transgenes as previously described (*45*). In this reporter line, GFP was inserted into the endogenous *CDKN1A* locus, while H2B-Cy5 served as a nuclear marker. Previously described MAFB^mTAGBFP^ (iPSC B), and HNF4A^YFP^ (iPSC C) reporter lines were also used in this study (*46*). All lines were cultured under the conditions described above and were regularly tested and confirmed negative for mycoplasma contamination.

### Directed differentiation to proxiloids

Differentiation was performed using a suspension-based protocol adapted from Chandrasekaran *et al*. (17). One day prior to differentiation (day −1), hiPSCs were dissociated with 1X TrypLE (ThermoFisher Scientific, cat.no 12563029) and seeded into AggreWell-800 plates (StemCell Technologies, cat.no. 34825) at 1.5 x 10^4^ cells per microwell in mTeSR^TM^ Plus (StemCell Technologies, cat.no. 100-0276) supplemented with 10 μM Y-27632 (Tocris, cat.no 1259). After 18-24 hours, embryoid bodies (EBs) were released according to the manufacturer’s instructions. For each differentiation, one well containing 1,500 EBs was transferred to five 60 x 15 mm Petri dishes (Greiner, cat.no. 628161), each containing 5 mL differentiation medium and placed on an orbital shaker set to 60 rpm (ThermoFisher Scientific, cat.no. 88881102).

The differentiation process consisted of three stages. During Stage 1 (primitive streak induction), EBs were transferred into “Medium 1,” which comprised a 1:1 mixture of DMEM without glucose (Gibco, cat.no. 11966-025) and Ham’s F12 (Gibco, cat.no. 21765029), supplemented with 2 mM GlutaMAX (ThermoFisher Scientific, cat.no. 35050061), 5 μg/mL insulin, 5 μg/mL transferrin, 5 ng/mL sodium selenite (Sigma-Aldrich, cat.no. I1884-1VL), 0.1% polyvinyl alcohol (Sigma-Aldrich, cat.no. P8136-250G), and 0.1% methylcellulose (Sigma-Aldrich, cat.no. M7027-100G). On day 0, Medium 1 was supplemented with 3 μM CHIR99021 (Tocris, cat.no 4423), 0.5 μM TTNPB (Merck Millipore, cat.no. T3757), and 10 μM Y-27632, and organoids were cultured for 42 h. Medium was then replaced with Medium 1 containing 0.5 μM TTNPB for an additional 30 h.

Stage 2 (proximal fate specification) began with the transition to “Medium 2,” which consisted of Medium 1 plus 10 ng/mL epidermal growth factor (Miltenyi Biotech, cat.no. 130-093-825) and 36 ng/mL hydrocortisone (Sigma-Aldrich, cat.no. H0888-1G). For this stage, the medium was further supplemented with 10 ng/mL human recombinant FGF9 (Biotechne R&D, cat.no. 273-F9-025) for 65 h to promote proximal tubule lineage commitment.

Stage 3 (maturation) commenced on day 6, when organoids were maintained in Medium 2 without FGF9. Medium was exchanged every 2–3 days until day 14–21, depending on the experimental endpoint.

### Directed differentiation to 2D PT-like cells

For matched 2D comparisons, hiPSCs were differentiated into PT-like cells using the previously described protocol of Chandrasekaran *et al*. (17). Briefly, hiPSCs were seeded at 3.5 x 10^4^ cells/cm^2^ and were subjected to directed differentiation toward the proximal tubule lineage using the same differentiation sequence and media described in the original protocol. Following differentiation, cells were used in gentamicin dose-response experiments at day 14 of differentiation. All matched 2D cultures were generated from the same hiPSC source and differentiation experiment as the corresponding proxiloid cultures.

### Radius quantification of proxiloids

Feret’s diameter was measured for ten organoids from hiPSC lines A, B, and C at day 0, day 3, and day 14 of differentiation using ImageJ (version 2.9.0/1.53t; National Institutes of Health, Bethesda, MD, USA). To assess morphological reproducibility, the coefficient of variation (CV) was determined from the descriptive statistics function in GraphPad Prism (version 10.6.1), calculated within individual differentiation batches and across independent batches.

### Bulk RNA-sequencing

For bulk RNA sequencing, hiPSC lines iPSC0028 and MAFB^mTAGBFP^ were differentiated in parallel to day 14. Approximately 300 proxiloids per replicate were pooled and lysed in RA1 lysis buffer (Macherey-Nagel, cat.no 740961.500). Total RNA was extracted using the NucleoSpin RNA kit (Macherey-Nagel, cat.no 740955.250) according to the manufacturer’s instructions and stored at −80°C. Three independent differentiations were performed per line. Samples were shipped to GenomeScan (Leiden, the Netherlands) for library preparation and sequencing. For etoposide-exposure experiments, iPSC0028-derived proxiloids were treated on day 14 with 30 µM etoposide in Medium 2 for 24h, alongside controls, after which ∼300 proxiloids per replicate were lysed in RA1 buffer. RNA was extracted using the Maxwell RSC simplyRNA kit (Promega, cat.no. AS1390). RNA integrity was confirmed (RIN ≥ 7). Libraries were prepared by GenomeScan and sequenced on an Illumina NovaSeq platform generating 150 bp paired-end reads. All samples met quality criteria (Q30 ≥ 80%) and were included in downstream analysis.

### Bulk RNA-sequencing analysis

Raw sequencing data from in-house experiments and a publicly available 2D proximal tubule-like cell dataset (ENA: PRJEB23622/ERP111777) were processed using a single unified pipeline in Miniconda. Adapters were detected and reads were trimmed, including a 15 bp 5’ trim of both mates. Reads were aligned to the GRCh38.p14 human reference genome (Ensemble v113) using STAR (2.7.11b) with default parameters. Gene-level quantification of exonic reads was performed with featureCounts (Rsubread, v.2.24.0) in paired-end mode, producing a raw count matrix. Ensembl gene identifiers were mapped to HGNC symbols using biomaRt (v2.66.0), and counts for identifiers sharing a gene symbol were summed. Ribosomal (RPL, RPS), mitochondrially encoded (MT-), and unannotated genes were removed. Genes with at least 10 counts in at least three samples were retained. Analyses were performed in R (v4.5.0) using DESeq2 (v1.50.2). Counts were normalized by the median-of-ratios method, and a variance-stabilizing transformation (VST) was applied for correlation analysis (Spearman’s ρ) and expression heatmaps. For transporter and off-target lineage heatmaps, absolute VST values were displayed, averaged per cell line. For proximal tubule signature analyses, a curated 164-gene PT signature derived from human kidney single-cell transcriptomic datasets reported by Humphreys and colleagues (*28*) was used. VST values were z-scored per gene across all samples so that each gene contributed equally regardless of absolute expression. Signature genes were further assigned to eight manually curated functional categories, and category scores were calculated as the mean z-scored expression of genes within each category; negative scores therefore indicate below-average relative expression. For the etoposide experiment, differential expression between treated and control samples was assessed with the Wald test, and log2 fold changes were shrunk using apeglm for visualization. P values were adjusted by the Benjamini-Hochberg method, and genes with an adjusted P value below 0.01 and an absolute log2 fold change greater than 1.5 were considered differentially expressed. Gene set enrichment analysis of Hallmark gene sets (MSigDB, via msigdbr) was performed on the log2 fold change-ranked gene list using fgsea with a minimum set size of 10 and a maximum of 500.

### Immunofluorescence

Organoids were fixed in ice-cold 4% methanol-free paraformaldehyde (ProSciTech, cat.no. C004) for 20 minutes at 4 °C, followed by three washes in 0.3% Triton X-100 in PBS. For immunostaining, organoids were washed three times at RT for 30 minutes per wash in PBS containing 0.3% Triton X-100 and 0.5% 1-Thioglycerol (washing buffer). Hereafter, organoids were incubated with primary antibodies for 3 days at 37°C on a platform-rotator in blocking buffer (2% BSA, 0.3% Triton X-100 in PBS). Samples were washed for 4 h on a platform-rotator at 37°C after which the washing buffer was replaced with fresh washing buffer for an additional washing step overnight. The next day, organoids were incubated with secondary antibodies and DAPI diluted in blocking buffer (1 μg/mL) for 2 days, followed by a 4 h wash in washing buffer. For optical clearing, organoids were incubated in Ce3D clearing solution (22% N-methylacetamide, 80% Histodenz (wt/vol), 0.1% (vol/vol) Triton X-100, and 0.5% 1-thioglycerol) overnight at room temperature before embedding in a silicon isolator cassette for imaging. Primary antibodies included: biotinylated LTL (Vector Laboratories, B-1325, 1:200), rabbit anti-LRP2 (Abcam, ab76969, 1:50), rabbit anti-CLDN10 (Biorbyt, orb48053, 1:50), mouse anti-AQP1 (Abcam, ab9566, 1:200), rabbit anti-SLC3A1 (Proteintech, 16343-I-AO, 1:100), rabbit anti-laminin (Sigma-Aldrich, L9393, 1:100), rabbit anti-cleaved caspase 3 (Cell Signaling, 9661S, 1:400), rabbit anti-WT1 (Abcam, ab89901, 1:400), rabbit anti-SLC12A1 (R&D, AF2605, 1:400), rabbit anti-SLC12A3 (Biorby, orb100812, 1:400), and mouse anti-MEIS1/2/3 (Active motif, 39795, 1:100) and mouse anti-MAP2 (ThermoFisher, MA5-12826, 1:300). Primary antibodies were detected with donkey anti-rabbit Alexa Fluor 568 (Thermo Fisher Scientific, cat.no. A10042), donkey anti-mouse 647 (Thermo Fisher Scientific, cat.no. A31571), and streptavidin Alexa Fluor 488 (Thermo Fisher Scientific, cat.no. S11223) diluted 1:2000 in blocking buffer. Imaging was performed on a Dragonfly spinning-disk confocal microscope (Andor Technology) and post-processing was performed using Imaris 9.5 software (Bitplane, Oxford Instruments)

### Transmission Electron Microscopy

For morphological analysis, proxiloids were fixed for 1.5 hours, at room temperature, with 2.5% glutaraldehyde and 2% paraformaldehyde (Electron Microscopy Sciences, cat. no. 16210 and 15710 respectively) in 0.1M sodium-cacodylate buffered solution, pH 7.4. Subsequently, the probes were rinsed 2x with 0.1M sodium cacodylate-buffered solution, and fixed for 1 hour on ice in 1% osmium tetroxide (Electron Microscopy Sciences, cat. no. 19152) and 1.5% potassium ferrocyanide (Merck, cat. no. 4984) in sodium cacodylate buffer. Samples were further washed twice with demineralized water, twice with 70% ethanol, then dehydrated overnight in 70% ethanol, followed by 80% ethanol (10 min), 90% ethanol (10 min), and 100% ethanol absolute (2 x 15 minutes; 1 x 30 minutes). Proxiloids were afterwards infiltrated with mixtures of 25%, 50%, and 75% epon LX-112 (Ladd Research) in anhydrous acetone, HPLC grade (J.T.

Baker), in steps of 30 minutes, followed by infiltration with pure epon for 90 minutes at room temperature, then 30 minutes at 60°C. Consequently, the probes were embedded in pure epon, mounted in BEEM capsules (Agar Scientific), and polymerized for 48 hours at 60°C. Ultrathin sections (100 nm) were cut using a diamond knife (Diatome) and mounted onto copper slot grids (Science Services), covered with formvar film and a 2.5 nm carbon layer. To enhance the contrast, sections were stained with an aqueous solution of 7% uranyl acetate for 20 minutes, followed by Reynold’s lead citrate for 10 minutes. Samples were analyzed at an acceleration voltage of 120 kV using a FEI Tecnai G2 12 TWIN electron microscope, equipped with a Gatan OneView camera. Image mosaics containing hundreds of images each, were recorded with binning 2, at 6500x magnification, corresponding to a 3.4 nm pixel size at the specimen level. Images were collected and mounted together using automated data acquisition and stitching software (*47*). These extensive digital images give overviews of entire structures, allowing zooming in to high detail for qualitative analysis. Virtual slides were analyzed and annotated using Aperio ImageScope viewing software version 12.4 (Leica Biosystems).

### P-gp and OAT transporter function assays

Transporter activity of P-glycoprotein (P-gp) and organic anion transporter 1 (OAT1) was assessed using calcein-AM efflux and fluorescein uptake, respectively. For both assays, organoids were collected and washed twice in HBSS, supplemented with 10 mM HEPES (HBSS–HEPES, StemCell Technologies, cat.no 37150) for the calcein-AM efflux assay. For the P-gp-mediated efflux assay, organoids were incubated for 1 h at 37 °C in HBSS–HEPES containing calcein-AM (0.25 µM, Invitrogen, cat.no R37601), either in the absence or presence of the P-gp inhibitor PSC833 (5 μM, Tocris, cat.no 4042). A dye-free condition was included to determine background fluorescence. Following incubation, organoids were washed twice in HBSS–HEPES and lysed in 0.1% Triton X-100 in HBSS–HEPES in black 96-well plates. Intracellular fluorescence was measured at 488/518 nm (excitation/emission), and P-gp transport activity was expressed as the ratio of fluorescence in PSC833-treated versus untreated organoids after background correction.

OAT transporter function was evaluated by quantifying intracellular fluorescein accumulation. Organoids were exposed for 10 minutes at 37 °C to fluorescein (1 μM, Sigma-Aldrich, cat.no F6377) prepared in HBSS, with or without the OAT inhibitor probenecid (500 μM, Sigma-Aldrich, cat.no P8761). Following exposure, organoids were washed three times with ice-cold HBSS and transferred to black, clear-bottom plates containing 0.1 M NaOH. After a 10 minutes incubation at room temperature under gentle agitation, lysates were analyzed by fluorescence detection at 490/518 nm. OAT-dependent uptake was determined by comparing fluorescein signal between exposed and probenecid-treated conditions after subtraction of background fluorescence.

### OCR measurement

Proxiloids were exposed to medium or gentamicin (4 mg/mL, Life Technologies, cat.no 15750060) for 24 or 48 hours, after which they were plated on Seahorse XFe96 Spheroid Microplates (Agilent, cat.no 102978-100) and incubated for 1 h in freshly made Seahorse XF basal medium (DMEM, Agilent, cat.no 103680-100) with 10 mM glucose (Agilent, cat.no 10-357-7100), 1 mM pyruvate (Agilent, cat.no 10-357-8100), and 2 mM glutamine (Agilent, cat.no 103579-100). After 45 minutes incubation without CO_2_, plates were loaded into a XFe96 extracellular flux analyzer (Seahorse Bioscience). Mitochondrial respiration was assessed with oligomycin (8 μM, ATP synthase inhibitor), FCCP (2.5 μM, an uncoupling agent that collapses the proton gradient), and a mixture of rotenone (2 μM, Complex I inhibitor from electron transport chain) and antimycin A (2 μM, Complex III inhibitor from electron transport chain). Compounds were added sequentially, and the oxygen consumption rate was measured in 3-minute periods with 3-minute mixing during each cycle. For normalization, each organoid was lysed individually in 100 µl RIPA buffer with protease inhibitor (1:100) for 10 minutes on ice and 25 µL of the lysate was used for BCA protein quantification (Pierce, ThermoFisher Scientific, cat.no A55865).

### P21 reporter line toxicity

For DNA damage reporter assays, 3 organoids at day 14 were transferred into each well of a glass-bottom 96 well plates and incubated with 0 μM, 1 μM, 20 μM, 30 μM, or 50 μM of etoposide (Sigma-Aldrich, E1383) in Medium 2 under standard culture conditions. Organoids were imaged every 6 h at 37 °C and 5% CO_2_, using a Nikon TiE2000 confocal microscope.

### Cell viability assay

Cell viability was measured using WST-8 assay (Sigma-Aldrich, cat.no 96992-500TESTS-F). After exposure to gentamicin, 10 μL of WST-8 reagent was added to the cell culture medium and cells were incubated for 1 h at 37 degrees. The absorbance was measured at 450 nm using a microplate reader. Data was normalized to untreated cells and presented as a relative percentage of viability of untreated cells. Triton X-100 at 10% was used as a positive control.

### Lactate Dehydrogenase (LDH) assay

The evaluation of cell membrane integrity upon exposure to gentamicin was evaluated by measuring the extracellular LDH activity. LDH is an intracellular enzyme which catalyzes NADH lactate to pyruvate. LDH is released in the supernatant from the cytosol upon cellular damage. LDH activity was measured using the Cytotoxicity LDH-glo assay (Promega, cat.no J2380), following the manufacturer’s protocol. Briefly, 50 μL of cell supernatant was added to a white opaque 96-well plate after 24 h exposure to a dose-range of gentamicin. 50 μL of assay buffer was added to a final volume of 100 μL per well, after which the plate was incubated for 30 minutes at room temperature. After 30 minutes, luminescence was measured in a microplate reader. Triton X-100 at 10% was used as a positive control and untreated cells as a negative control. LDH activity was expressed as percentage of the positive control.

### Dose-response curve fitting and statistical analysis

Dose-response relationships were analyzed in R using the drc package (*48*). For each culture format (2D and proxiloids), responses were fitted with a four-parameter log-logistic model (4PL). For WST-8 viability measurements, the upper asymptote was constrained to 100% to reflect normalization to vehicle controls. For LDH release measurements, the lower asymptote was constrained to 0%, while all remaining parameters were estimated freely. To enable log-scale visualization and model fitting, the vehicle condition (0 µg/mL gentamicin) was assigned a nominal concentration of 1 µg/mL; response values were not altered. Data are presented as mean ± SEM (n = 3 biological replicates per concentration) together with the fitted 4PL dose-response curves. Effective concentrations (EC50/IC50) were estimated from the fitted models. Differences in sensitivity between 2D cultures and proxiloids were assessed by using ratio-based parameter comparison tests between fitted curves. Model-level differences between shared-curve and separate-curve fits were evaluated using an extra sum-of-squares F-test.

### Mitotracker and TMRM live staining

After 3 days of treatment, the manufacturer’s protocol for Mitotracker and TMRM (Thermofisher, cat.no M7514 and cat.no I34361 respectively) staining was followed. In short, proxiloids were live stained with a concentration of 100 nM MitoTracker and 0.1 µM TMRM in Medium 2. Proxiloids were incubated for 40 minutes at 37 degrees. After incubation, cells were washed 3x with Medium 2 and imaged using the Airyscan NIR M980, 40x zoom. Mitochondrial network morphology was quantified in ImageJ using the Mitochondria Analyzer plugin. Briefly, mitochondrial masks were generated from MitoTracker images and network connectivity was assessed by calculating the longest shortest path, which was used as a measure of mitochondrial network integrity and branching.

### Mitochondrial DNA copy number quantification

Total DNA was extracted from proxiloids after seven days of compound exposure using Qiagen DNA extraction kit (cat.no 69504) according to the manufacturer’s instructions. For each condition, 5 proxiloids were pooled per sample and 3 biological replicates were analyzed. DNA concentration and purity were measured by Nanodrop, and DNA was diluted to 10 ng/μL in nuclease-free water. Mitochondrial DNA copy number was determined by quantitative PCR using primers targeting the mitochondrial gene MT-ND1 and the nuclear single-copy reference gene B2M. Reactions were performed in 10 µL containing 1x iQ SYBR Green Supermix (Bio-Rad, cat.no 1708886), 500 nM of each primer, and 10 ng template DNA. Amplification was carried out the CFX Connect Real-Time System (Bio-Rad) under the following conditions: denaturation at 95°C for 3 minutes, followed by 40x cycles of 95°C for 10 sec, 60°C for 30 sec, and 72°C for 30 sec. All samples were run in triplicates. Primer sequences are shown in Table 1. Relative mitochondrial DNA copy number was calculated using the ΔCt method. For each sample, ΔCt was defined as Ct(MT-ND1) − Ct(B2M), and the relative MT-ND1 to B2M ratio was calculated as 2^(−ΔCt). Results are expressed as mtDNA copy number (MT-ND1/B2M ratio). Statistical comparisons between control and treated conditions were performed using one-way ANOVA with Dunnett’s post hoc test, with significance defined as p < 0.05.

**Table 1.**
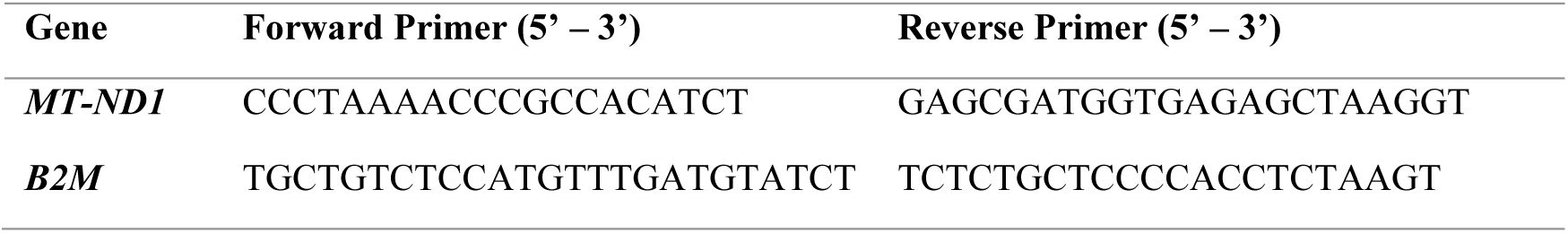
Primer sets used for quantification by Real-time PCR.

### Cellular ATP quantification

Cellular ATP content was measured as a readout of metabolic viability using the CellTiter-Glo 3D Cell viability Assay (Promega, cat.no G9682), selected for its enhanced lysis capacity in three-dimensional structures relative to the standard formulation. Adefovir-exposed proxiloids were placed in white-walled 96 well plates after seven days of exposure. Each well contained 100 µL of CellTiter-Glo reagent and one proxiloid. Contents were mixed in the dark on a plateshaker for 5 minutes to induce cell lysis. The plate was then allowed to incubate at RT for an additional 25 minutes to stabilize the luminescent signal, and luminescence was recorded using an integration time of 0.25 second per well.

Luminescence values were background-corrected against blank wells containing reagent without organoids and normalized to vehicle-treated controls. Values were expressed as percentage of control ATP content. All samples were run using 6 technical replicates and each experiment has been performed in biological triplicates. Statistical comparisons between control and treated conditions were performed using one-way ANOVA with Dunnett’s post hoc test, with significance defined as p < 0.05.

### Statistical analysis

Every experiment was performed in at least three biological replicates, including at least three technical replicates each, unless specified otherwise. Results are presented as mean ± the standard deviation (SD) or standard error of the mean (SEM), as indicated, and *P* values < 0.05 were considered statistically significant. Numeric data was analyzed using GraphPad 9.0 (GraphPad software, La Jolla, CA, USA). For statistical analysis, one-way ANOVA was performed followed by Dunnett’s post-hoc analysis unless specified otherwise. Image analysis was performed with ImageJ (ImageJ software version 2.9.0/1.53t, National Institutes of Health, Bethesda, MD, USA).

### Ethics statement

All experiments involving human-derived material used previously established cell lines or previously collected human tissue. The parental iPSC0028 line was obtained commercially from Sigma-Aldrich. The CDKN1A^GFP^-H2B^Cy5^ reporter line was generated from the iPSC0028-HC3X background by CRISPR/Cas9-mediated genome engineering in the laboratory of Bob van de Water (Leiden University). The HNF4A^YFP^ and MAFB^mTagBFP^ reporter lines were generated from commercially obtained human cells, as described previously (*46*). The original donor materials for these commercially sourced cells were obtained by the suppliers under appropriate informed-consent procedures.

Adult human kidney tissue was not newly collected as part of this study. TEM images of adult human kidney tissue shown for comparison were generated from previously collected cortical tissue. This kidney was not accepted for transplantation and tissue was obtained with research consent (*49*). The adult human kidney images used in the present study are distinct images from those reported in the original publication and are used here for comparative ultrastructural analysis.

## Acknowledgments

We thank Melissa Little (Murdoch Children’s Research institute, Melbourne, Australia) for providing the MAFB^mTagBFP^ and HNF4A^YFP^ cell lines. We gratefully acknowledge Manon Zuurmond (LUMC, Leiden, the Netherlands) for assistance with the schematic figures, and Anja Wilmes (VU Amsterdam, the Netherlands) for the advice on the 2D cultures. We thank Elpida Lymperi for her help and troubleshooting. We also acknowledge the technical support provided by Johan W.A. Verspuij and thank the Light and Electron Microscopy Facility (LUMC, Leiden, the Netherlands) for their technical assistance and maintenance of the microscopes.

## Author contributions

Conceptualization, S.B., H.S.S., M.H., B.v.d.W, C.W.v.d.B., T.J.R.; methodology, S.B., H.S.S., M.H.; software, S.B., R.H.S., M.H.; validation, S.B., D.N.v.H., H.S.S; formal analysis, S.B., M.H.; investigation, S.B., M.C.A., D.N.v.H., R.v.N., R.F.V.; resources, H.S.S, T.J.R., M.H, B.v.d.W; data curation, S.B.; writing – original draft, S.B.; writing – review & editing, S.B., H.S.S, M.H, B.v.d.W, T.J.R; visualization, S.B.; supervision, H.S.S, M.H, T.J.R; project administration, H.S.S, M.H, T.J.R; funding acquisition, H.S.S, T.J.R. All authors have read and agreed to the published version of the manuscript.

## Data, code, and materials availability

All data needed to evaluate the conclusions in the paper are present in the paper and/or the Supplementary Materials. Raw bulk RNA sequencing data generated during the current study is available at the European Nucleotide Archive (ENA) at the EMBL-EBI under the accession numbers XXX (datasets named…). The 2D dataset was accessed under accession numbers PRJEB29471/ERP111777. The code used in this study is available at GitHub (LINK).

---

**Fig. S1.**
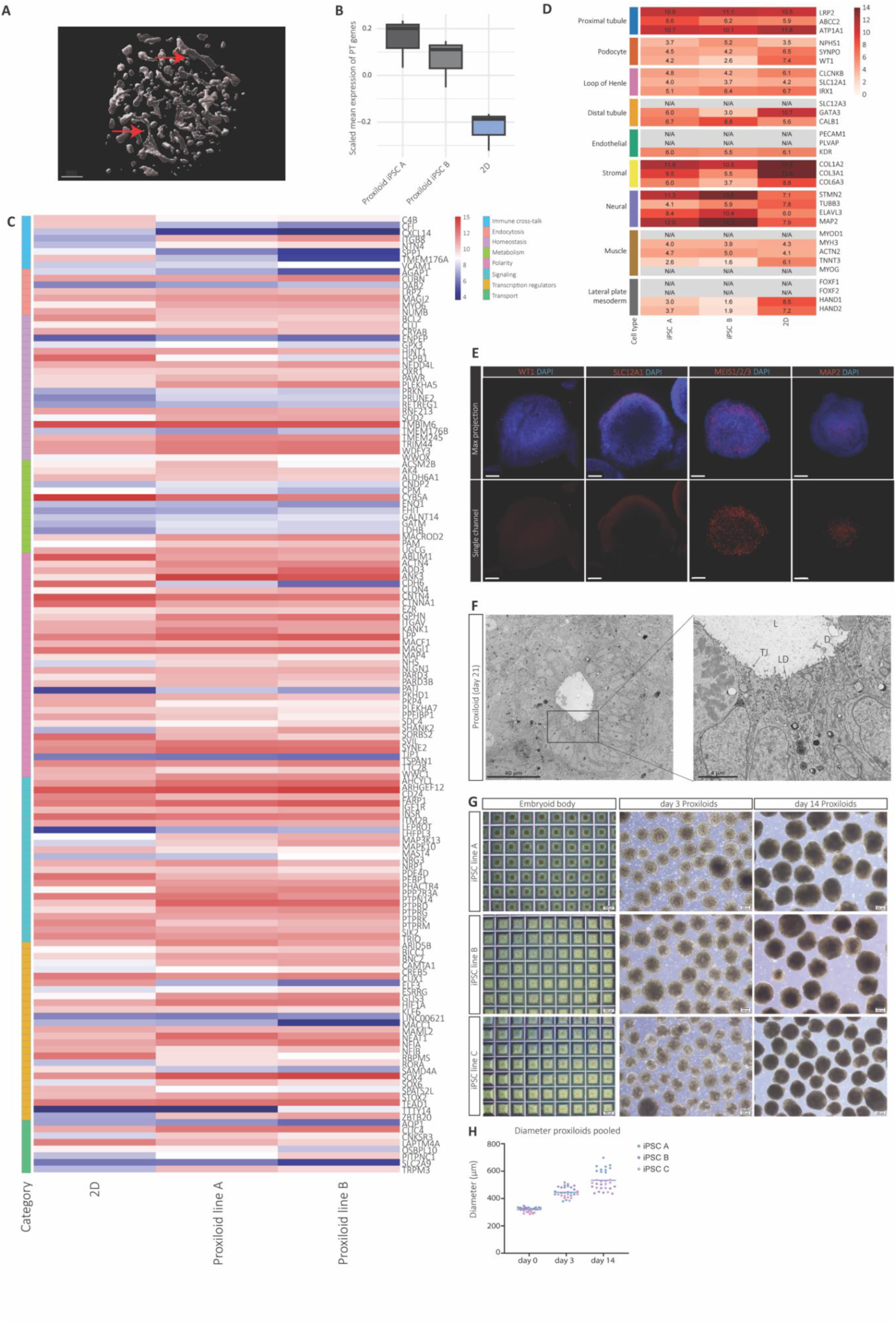
Proxiloid architecture and identity. **(A)** Three-dimensional reconstruction of proxiloids showing an interconnected network of lumenized epithelial tubules. Arrows indicate representative elongated tubular segments. Scale bar, 100 µm. **(B)** PT signature score (gene-wise scaled expression of a curated 164-gene PT signature) comparing proxiloids and matched 2D monolayers from two hiPSC lines (n = 3 differentiations per line). **(C)** Heatmap showing variance-stabilized expression values of the curated PT signature genes in 2D cultures and proxiloids from two hiPSC lines. Genes are grouped by functional category. **(D)** Heatmap showing variance-stabilized expression of representative lineage markers across two hiPSC lines. PT markers are enriched, while markers of other nephron segments are low or absent. Stromal-associated genes (*COL1A2*, *COL3A1*, *COL6A3*) and neural-associated genes (*STMN2*, *ELAVL3*, *MAP2*) are detectable. Genes below the expression threshold are indicated as N/A. **(E)** Confocal immunofluorescence for WT1, SLC12A1, MEIS1/2/3, and MAP2 (red); DAPI labels nuclei (blue). Maximum intensity projections (top) and corresponding single-channel images (bottom) are shown. Scale bars, 100 µm. **(F)** Transmission electron microscopy (TEM) of day 21 proxiloids showing expanded luminal structures and organized epithelial architecture. Left: low-magnification overview of a proxiloid with a central lumen. Right: higher-magnification view highlighting epithelial morphology. L, lumen; TJ, tight junction; LD, lipid droplet; D, desmosome. Scale bars, 40 µm (left) and 4 µm (right). **(G)** Brightfield images of proxiloids derived from three independent hiPSC lines at day −1 (scale bar, 500 µm), and days 3 and 14 (scale bars, 200 µm). **(H)** Proxiloid diameter distributions at day 0, day 3, and day 14 across independent differentiations and hiPSC lines. Distributions were used for coefficient of variation (CV) analysis to assess intra- and inter-batch reproducibility.

## Movie S1

**Movie S1. 3D reconstruction of proxiloid tubular network (LRP2, red). Related to Fig. 1 and Fig. S1.**

## Data S1. (separate file)

**Data S1. Manually curated list of PT identity genes, based on single cell data** (***28***).

